# Biomimetic multi-channel microstimulation of somatosensory cortex conveys high resolution force feedback for bionic hands

**DOI:** 10.1101/2023.02.18.528972

**Authors:** Charles M. Greenspon, Giacomo Valle, Taylor G. Hobbs, Ceci Verbaarschot, Thierri Callier, Elizaveta V. Okorokova, Natalya D. Shelchkova, Anton R. Sobinov, Patrick M. Jordan, Jeffrey M. Weiss, Emily E. Fitzgerald, Dillan Prasad, Ashley van Driesche, Ray C. Lee, David Satzer, Jorge Gonzalez-Martinez, Peter C. Warnke, Lee E. Miller, Michael L. Boninger, Jennifer L. Collinger, Robert A. Gaunt, John E. Downey, Nicholas G. Hatsopoulos, Sliman J. Bensmaia

## Abstract

Manual interactions with objects are supported by tactile signals from the hand. This tactile feedback can be restored in brain-controlled bionic hands via intracortical microstimulation (ICMS) of somatosensory cortex (S1). In ICMS-based tactile feedback, contact force can be signaled by modulating the stimulation intensity based on the output of force sensors on the bionic hand, which in turn modulates the perceived magnitude of the sensation. In the present study, we gauged the dynamic range and precision of ICMS-based force feedback in three human participants implanted with arrays of microelectrodes in S1. To this end, we measured the increases in sensation magnitude resulting from increases in ICMS amplitude and participant’s ability to distinguish between different intensity levels. We then assessed whether we could improve the fidelity of this feedback by implementing “biomimetic” ICMS-trains, designed to evoke patterns of neuronal activity that more closely mimic those in natural touch, and by delivering ICMS through multiple channels at once. We found that multi-channel biomimetic ICMS gives rise to stronger and more distinguishable sensations than does its single-channel counterpart. Finally, we implemented biomimetic multi-channel feedback in a bionic hand and had the participant perform a compliance discrimination task. We found that biomimetic multi-channel tactile feedback yielded improved discrimination over its single-channel linear counterpart. We conclude that multi-channel biomimetic ICMS conveys finely graded force feedback that more closely approximates the sensitivity conferred by natural touch.

## Introduction

Manual interactions with objects rely critically on tactile signals from the hand, as evidenced by the deficits incurred when these signals are lost or eliminated *(1–3)*. With this in mind, efforts are under way to provide brain-controlled bionic hands with tactile feedback via intracortical microstimulation (ICMS) of somatosensory cortex (S1) *(4, 5)*, which has been shown to evoke vivid tactile percepts *(6, 7)* and improve control of prosthetic hands *(8)*. To be useful, tactile feedback needs to convey information about contact events, including information about contact location and force. Information about location can be intuitively conveyed by matching force sensors on the bionic hand with somatotopically appropriate electrodes in S1 *(6, 8, 9)*. For example, a force sensor on the index fingertip of the bionic hand drives stimulation of electrodes located in the index representation of S1, thereby evoking a sensation experienced on the index finger. Information about contact force can be conveyed by modulating ICMS amplitude according to the output of the sensor, where higher stimulation amplitudes give rise to more intense touch sensations, paralleling the sensory correlates of increases in force on the skin *(6, 9, 10)*.

The objective of the present study was to examine the precision and accuracy of ICMS-based tactile feedback about contact force in three human participants implanted with arrays of microelectrodes in S1. To this end, we first characterized the increases in sensation magnitude resulting from increases in ICMS amplitude (cf. ref *(6)*) and gauged the intensity of these percepts against tactile benchmarks. We found that the intensity of ICMS-evoked sensations was highly electrode-dependent and often faint. Furthermore, only a few discriminable levels of intensity could be achieved using standard force feedback, which consists of linearly modulating ICMS amplitude according to the applied force. Seeking to improve the range and precision of ICMS-based force feedback, we implemented “biomimetic” ICMS-trains, which, by emphasizing contact transients and de-emphasizing maintained contact, evoke patterns of neuronal activity that more closely mimic those in natural touch *(11, 12)*. We found that biomimetic ICMS yielded higher resolution force feedback than did its linear counterpart, even though the total charge of biomimetic ICMS spanned a narrower range. Next, we investigated whether we could further improve force feedback by delivering biomimetic ICMS through multiple electrodes simultaneously. We found that multi-channel biomimetic ICMS gives rise to stronger and more distinguishable sensations than does its single-channel counterpart, thus enabling precise force feedback over a wider range of forces. Biomimetic multi-channel ICMS more closely approximates the sensitivity conferred by natural touch than the other feedback regimes tested.

## Results

Three participants with cervical spinal cord injury were each implanted with four electrode arrays, two in the arm and hand representation of motor cortex and two in the hand representation in Brodmann’s area 1 of S1 (Figure 1A). In all three participants, stimulation through electrodes in S1 evoked sensations experienced on the contralateral hand, following the expected somatotopic organization (Figure 1B for C1, Supplementary Figure 1A for P2 and Supplementary Figure 1B for P3, also see *(13)*).

**Figure 1.**
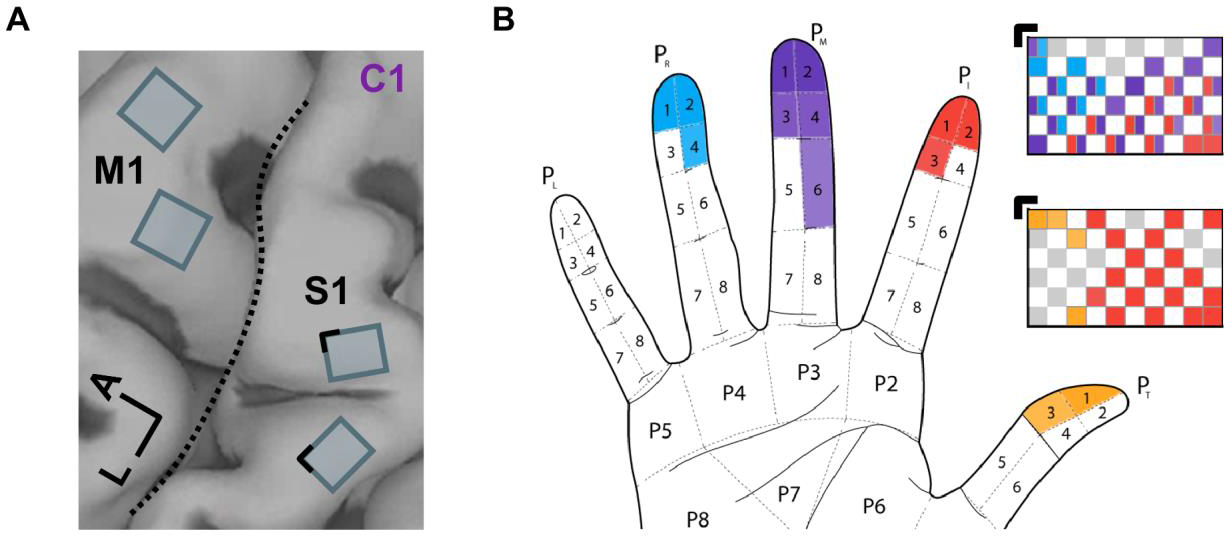
ICMS of S1 evokes tactile percepts whose location follows the expected somatotopic organization. **A|** Four Utah arrays (Blackrock Neurotech, Inc.) were implanted in participant C1, two of which were placed in the hand representation of S1, based on localization with fMRI. L: Lateral. A: Anterior. **B|** Locations of projected fields – the location on the hand where sensations are experienced – for each S1 channel for participant C1. The top array is medial, bottom one lateral. Colors denote the location of the projected field. Gray squares denote electrodes that evoked sensations on the dorsum of the hand, and white squares denote unwired electrodes. Black corners indicate alignment. See Supplementary Figure 1 for information about the implants of the other two participants (P2 and P3).

### The dynamic range of ICMS-evoked sensations is electrode-dependent

In natural touch, increases in contact force lead to increases in the firing rate of active neurons and in the recruitment of additional neurons in a somatotopically determined region of S1 *(11)*. Increasing the ICMS amplitude leads to the same qualitative pattern of neuronal activation *(14)* and to increases in the perceived magnitude of the evoked sensation *(6–8)*, analogous to the sensory correlates of force increases. With this in mind, we gauged the consistency of this relationship across participants and stimulating electrodes. The three participants rated the perceived magnitude of a sensation on a numerical scale of their choosing *(15)*. The ratings were then normalized by the mean rating for each electrode and their relationship with ICMS amplitude was characterized. On every electrode tested, perceived magnitude increased approximately linearly with ICMS amplitude, replicating previous findings (Figure 2A). The median correlation between amplitude and intensity rating was 0.97, with all but two correlations (both from participant P3) above 0.9 (range: 0.47 to 0.99). The relationship between ICMS amplitude and perceived magnitude is thus robust.

**Figure 2.**
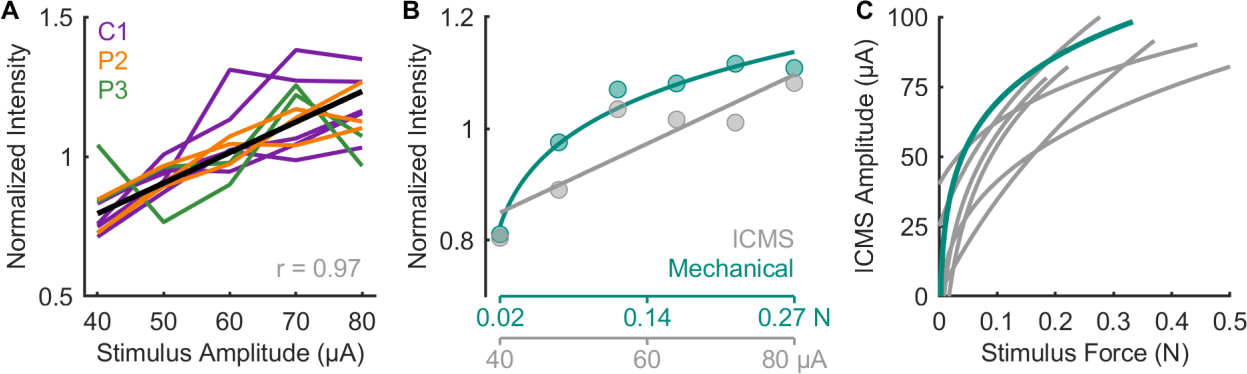
Perceived intensity increases linearly with ICMS amplitude. **A|** Normalized magnitude ratings following ICMS through single channels for 3 participants. Each line denotes ratings for one channel, different colors denote different participants. The thick black line denotes the mean across channels and participants. **B|** Normalized ratings when ICMS and mechanical stimuli are interleaved for one electrode and indentations delivered to the location on the skin corresponding to the PF of that electrode. **C|** Equal intensity contours for ICMS-evoked and mechanically evoked sensations. Each line represents the contour derived from the ratings from one stimulating channel. The teal line corresponds to the contour of the channel shown in panel B. While perceived magnitude increases with amplitude on all channels, the magnitude of the sensations varies widely across channels. Data from panels B and C are from participant C1 only.

The limitation of the magnitude estimation approach, however, is that ratings are (by virtue of the paradigm) participant-specific *(15, 16)*, so they cannot be benchmarked to natural touch. Furthermore, ratings are normalized within each block or session so ratings cannot be compared across sessions. To overcome these limitations, we leveraged the fact that participant C1’s tactile sensation on the hand is on par with that of able-bodied controls (Supplementary Figure 2A, B). Participant C1 judged the magnitude of the sensations evoked by ICMS trains and by skin indentations (delivered to a location matching the projected field of the stimulating electrode) of varying force, with the two types of stimuli interleaved randomly within each experimental block. We could then directly compare the magnitude of electrically and mechanically evoked sensations (Figure 2B) because both stimulus types were rated on the same scale. Furthermore, assuming the perceived magnitude of the tactile stimuli remained constant across sessions, we could also compare the perceived magnitude of ICMS delivered through different electrodes on different sessions. From these combined ICMS and touch sessions, we constructed equal intensity contours for the two stimulus types (Figure 2C), allowing us to determine the ICMS amplitude required to evoke a sensation whose intensity corresponds to a given force and vice-versa. First, we found that perceived magnitude increased approximately linearly with ICMS amplitude but as a (decelerating) power function of mechanical amplitude *(17)* (Supplementary Figure 2C). Accordingly, the equal intensity contours also followed a power law, reflecting the non-linearity in the mechanically evoked sensations (Figure 2C). Furthermore, while the magnitude of ICMS-evoked sensations increased with amplitude on all electrodes, the intensity range of the sensations varied widely across electrodes: some electrodes could only evoke weak sensations (< 0.25 N, the weight of a USB stick), whereas others could evoke sensations commensurate with moderate forces (∼ 0.5 N, the weight of an egg). Given that contact forces often exceed 1 N during object interactions *(18, 19)*, the dynamic range of ICMS-evoked sensations, constrained by the maximum safe stimulation level (100 μA, *(6, 20)*), is generally narrow.

### The discriminability of evoked percepts is electrode-dependent

Having established that the mapping between ICMS amplitude and perceived contact force was electrode dependent, we investigated how sensitive participants were to changes in ICMS amplitude. To this end, we had participants discriminate flat ICMS trains that varied in amplitude. On each trial, the participant reported which of two sequentially presented stimuli was more intense. The amplitude of the standard (or reference) stimulus was 60 μA and that of the comparison varied between 40 and 80 μA. From the behavioral performance, (Figure 3A), we computed, as an index of sensitivity for each electrode, the Just Noticeable Difference (JND), which denotes the change in stimulus amplitude that would yield a criterion level of performance (75% correct). While the median JND was around 13.5 μA (Figure 3B), consistent with previous results in humans and monkeys *(6, 7, 9, 10)*, JNDs varied widely across electrodes and participants (interquartile range: 8.5 to 22.9 μA).

**Figure 3.**
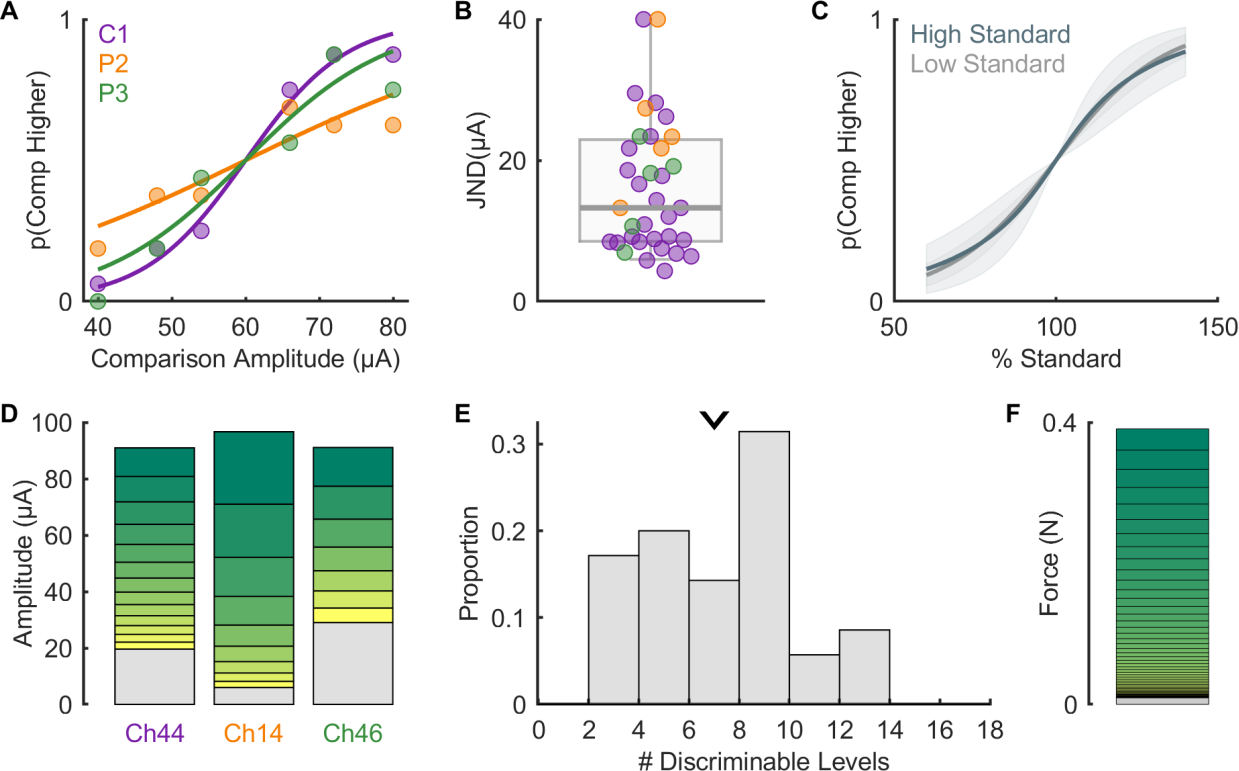
Stimulus discriminability is electrode dependent. **A|** Example psychometric functions from each participant – note that the curve for each participant is not representative of overall performance. **B|** JND computed from psychometric functions for all electrodes across participants (n = 35) – thick grey line denotes median (13.5 μA). Note that values above 40 μA are set to 40 μA for graphical purposes (n=2). **C|** Mean psychometric function (n = 8) when ICMS amplitude discrimination was performed with a high and a low standard. Comparison amplitudes are expressed as a percentage of the standard. The overlap between the two psychometric functions is consistent with Weber’s law, but not all the electrodes followed this relationship (see Supplementary Figure 3). **D|** Diagram showing the estimated discriminable levels plots for the electrodes shown in panel A. The grey section at the bottom of each bar indicates the subthreshold range for that electrode while the height of each subsequent bar is determined by the JND (following Weber’s law), with the maximum amplitude capped at 100 μA. **E|** The number of discriminable levels across all electrode and participants. Arrow indicates median number of discriminable levels (7). **F|** Estimated discriminable levels for tactile stimulation in the same approximate range of forces (0-0.4N).

To assess whether the discrimination of ICMS amplitude follows Weber’s law – with JNDs increasing in proportion to the standard amplitude, we tested amplitude discrimination at two different standards on a subset of electrodes. We found that, on average, JNDs followed Weber’s law, but the tendency to do so varied widely across electrodes (Figure 3C, Supplementary Figure 3). Having measured the detection threshold and the JND for each electrode, we computed the number of discriminable levels (Figure 3D), defined as the number of discrete, discriminable intensity levels between detection threshold and maximum safe amplitude (100 μA), assuming a constant Weber fraction (the ratio of JND to standard amplitude). We found that the number of discriminable steps ranged from 2 to 14, with a median of 7 (Figure 3E), implying that the force feedback conveyed by flat ICMS is coarse, allowing for only a handful of discriminable levels within the safe stimulation range. Electrodes with lower detection thresholds tended to have higher JNDs (r = −0.51, p < 0.01, Supplementary Figure 4A), suggesting that JNDs do not simply reflect overall sensitivity to ICMS. Electrodes with more discriminable levels also tended to evoke the most intense sensations (r = 0.87, p = 0.02, Supplementary Figure 4B), implying that the JND for each electrode corresponds to a change in intensity that is approximately consistent across electrodes, a phenomenon that is non-trivial given that magnitude functions are not systematically predictable from JNDs in natural perception *(21, 22)*. As a benchmark, the native touch of both participant C1 and able-bodied controls (n = 5) yielded around 45-50 discriminable levels over this span of forces, an order of magnitude more than what could be achieved with single-channel ICMS over the safe range of amplitudes (Figure 3F). Natural touch thus confers far greater sensitivity to force variations than does single-channel linear ICMS.

### Biomimetic feedback confers greater sensitivity to changes in force

In the experiments described above, ICMS consisted of pulse trains of constant amplitude (flat trains). Such pulse trains typically evoke an abrupt rise in the activation of neurons around the electrode, followed by a slow decrease *(23)*. In contrast, interactions with objects evoke phasic responses at the onset and offset of contact and much weaker responses (< 10%) during maintained contact, a property inherited from the periphery (11). In studies in amputees with electrical stimulation of the peripheral nerves, tactile feedback that features greater sensitivity to contact transients (thereby mimicking natural touch) has been shown to confer greater dexterity to myoelectric bionic hands *(24–26)*. With this in mind, we assessed participants’ ability to discriminate biomimetic ICMS trains, designed to mimic the S1 response to the onset, maintenance, and termination of contact (Figure 4A). We compared the participants’ performance with biomimetic trains to that with trains designed to track the forces linearly, which largely resembled flat trains with short on- and off-ramps. For this comparison, the biomimetic and linear trains were matched in peak amplitude, so the charge delivered in a biomimetic train was less than that in a matched linear one (69 ± 7 % on average, see Methods). We found that JNDs for biomimetic ICMS were systematically lower than were their linear counterparts (Figure 4, medians: 9.7 and 16.6 μA; IQRs: 7.7 to 16.1 μA and 11.5 to 23.5 μA, respectively, Wilcoxon signed-rank test, n = 22, z = 3.1, p < 0.01). The improved sensitivity to biomimetic ICMS is especially surprising given that biomimetic trains contained less charge. For the set of electrodes tested in this comparison, the median number of discriminable levels increased from 8 to 11 with biomimetic sensory feedback (Figure 4C, Wilcoxon rank sum test: Z = 3.21, p < 0.01). Biomimetic ICMS trains thus provide higher resolution force feedback, an advantage that holds true when expressed in terms of charge rather than peak amplitude (Supplementary Figure 5A,B) and does not simply reflect differences in perceived intensity (Supplementary Figure 5C).

**Figure 4.**
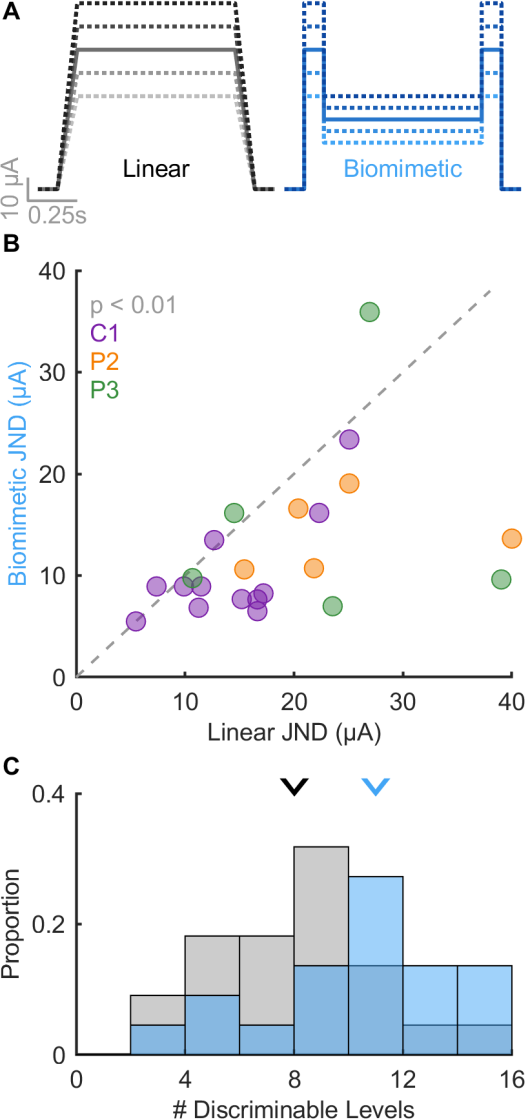
Biomimetic stimuli are more discriminable than linear stimuli. **A|** Example idealized stimulus profiles for linear (where stimulus amplitude scales with force) and biomimetic (where force transients are emphasized) stimuli. **B|** JNDs are reduced (sensitivity enhanced) with biomimetic stimuli versus linear ones. **C|** The distribution of the number of discriminable levels computed from JNDs with linear (gray) or biomimetic (blue) stimuli. The number of levels increases with biomimetic stimuli versus linear ones (median = 11 vs. 8, respectively).

### Multi-channel feedback confers greater dynamic range and change sensitivity

Having characterized the dynamic range and resolution of ICMS-based force feedback delivered through single-channel at a time, we next examined the magnitude of sensations evoked when ICMS was delivered through multiple electrodes simultaneously. For these experiments, we selected groups of 4 electrodes (quads) with overlapping or adjacent projected fields (the hand regions over which the sensations were experienced) (Supplementary Figure 6A). That is, when stimulation was delivered through each of the four electrodes individually, the sensation was experienced on an overlapping patch of skin or, in one case, on adjacent digits *(13)*. In these experiments, ICMS was always biomimetic (Figure 4A). First, we assessed whether multi-channel stimulation increased the dynamic range of the evoked sensations by comparing magnitude estimates of intensity with (biomimetic) single-vs. multi-channel stimulation (3 sets of 4 electrodes in participant C1). We found that multi-channel stimulation evoked systematically more intense sensations than did single-channel stimulation when equating the current delivered through each electrode individually (so quad stimulation delivered four times more total current than did its matched single-electrode counterpart, Figure 5A). Nonetheless, within the safe range of ICMS amplitudes, multi-channel ICMS allowed for a much wider dynamic range than did any electrode in isolation (more than twice the average electrode, Figure 5B). Indeed, the peak equivalent force reached 2 N, approximately the weight of a mobile phone. Furthermore, the multi-channel ICMS amplitude intensity function still followed a linear relationship with amplitude (Supplementary Figure 6B), so this linear relationship is not an artifact of testing a narrow range of intensities *(21)*.

**Figure 5.**
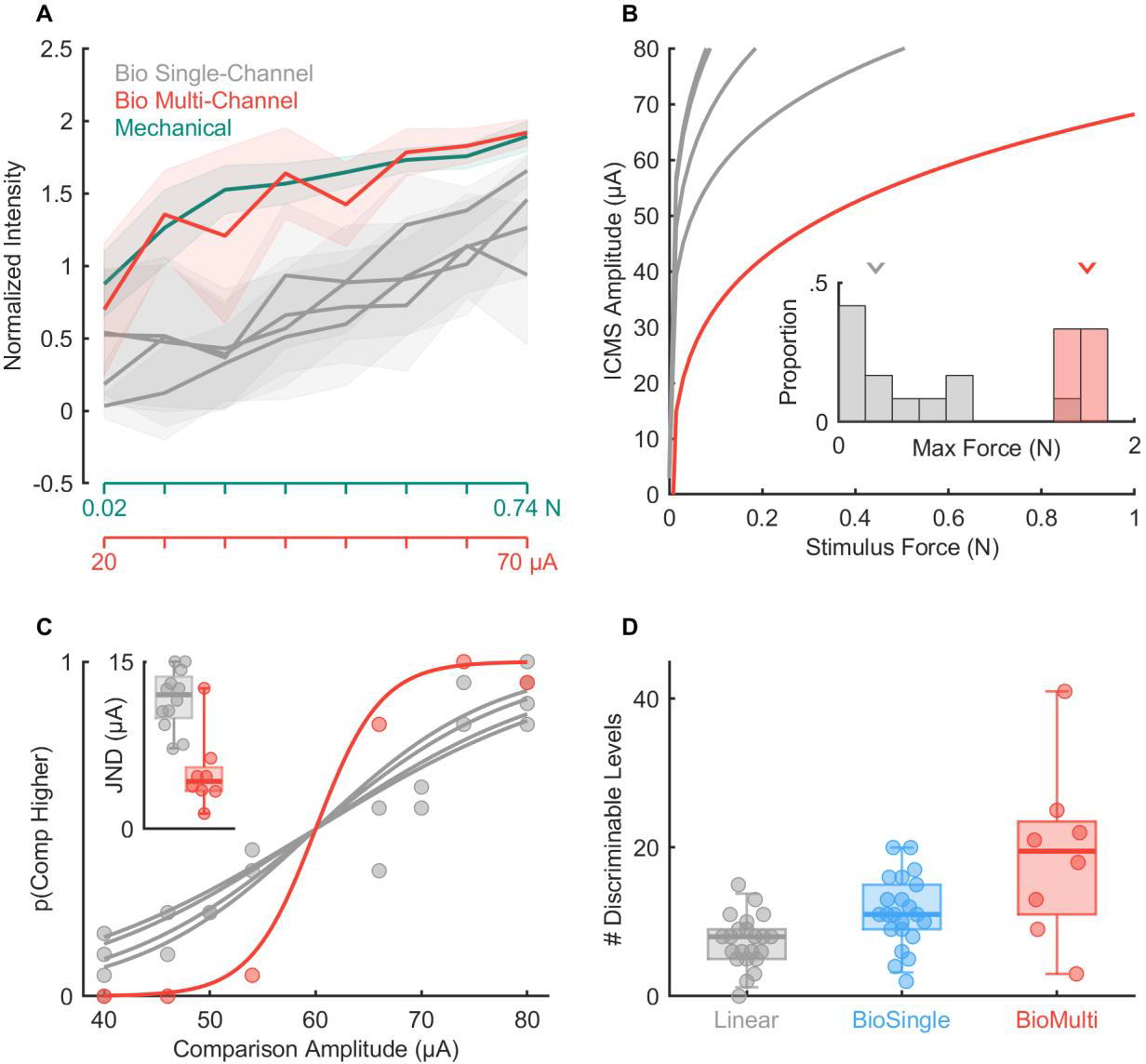
Multi-channel biomimetic stimulation produces more intense and more discriminable percepts. **A|** Normalized magnitude ratings when ICMS is delivered through electrodes individually or simultaneously, interleaved with mechanical stimuli for an example quad. All ICMS trains were biomimetic (cf. Figure 4A). Lines denote the mean while shaded areas denote the standard deviation. **B|** Equal-intensity contours for single-channel (gray) and multi-channel (red) ICMS for an example quad. Inset: Maximum achievable force for single or multi-channel stimulation across all tested quads, extrapolating fits to 100 μA. **C|** Psychometric functions for one quad of electrodes with the performance of the individual electrodes shown in gray and that of the quad in red. Inset: JNDs for all single electrodes and quads tested. The median JND decreased from 12 to 4.3 μA with multi-channel ICMS (Wilcoxon rank sum test: Z = −3.05, p < 0.01). **D|** Estimated number of discriminable levels with single-channel ICMS (linear or biomimetic stimuli, median = 8 and 11, cf. Figure 4C) and multi-channel biomimetic ICMS (median = 19.5). Panels A & B show results from participant C1 while panels C & D show results from participants C1 and P2.

Next, we examined the discriminability of multi-channel biomimetic ICMS trains that varied in amplitude (n = 8 quads, 5 from participant C1 and 3 from participant P2). We found that multi-channel ICMS – with identical stimulation delivered through 4 channels – yielded substantially lower JNDs than did its single-channel counterpart when expressed in terms of the amplitude on each electrode (Figure 5C), mirroring the lower variability in the magnitude estimates of intensity for multi-channel stimulation compared to its single channel counterpart (Supplementary Figure 6C). Combined, the wider dynamic range and higher resolution yielded an increase in the median number of discriminable levels from 11 to 19.5 (Figure 5D). Thus, biomimetic multi-channel force feedback can yield more precise force feedback over a wider dynamic range than standard force feedback through a single electrode. Indeed, while JNDs expressed in terms of charge were equivalent or higher for multi-channel than single-channel stimulation (Supplementary Figure 6D), the bottleneck in ICMS-based feedback is the charge delivered on any given electrode, which has been identified as the primary determinant of stimulation-induced neuronal damage *(27)*. Multi-channel ICMS circumvents this limitation by distributing the charge, thereby increasing both the range and precision of the resulting sensations to a level that more closely approximates natural touch (∼20 vs. 45-50 discriminable steps).

### Multi-channel biomimetic ICMS provides more intuitive force feedback in a closed-loop task

Having established that that multi-channel biomimetic feedback leads to improved sensitivity in passive discrimination experiments, we implemented this stimulation paradigm in a bionic hand (Ability Hand, Psyonic) and had participant C1 use it to perform a closed-loop task that required force perception, namely compliance discrimination. In brief, we trained a long short-term memory (LSTM) network to decode hand aperture from neural signals harnessed from M1. Real-time feedback from the sensors on the hand about the forces exerted on the object was conveyed via linear feedback or biomimetic multi-channel feedback, with different feedback conditions tested in separate experimental blocks. For linear feedback, ICMS amplitude was proportional to time-varying sensor output; for biomimetic feedback, ICMS amplitude was proportional to sensor output and its derivative. On each trial, the (blinded) participant grasped two objects – foam pucks of compliance 3 lb/ft^3^ and 8 lb/ft^3^ (FlexFoam, SmoothOn), presented in random order – and reported which of the two was firmer (Supplementary Figure 7). Consistent with results from the passive discrimination experiments, we found that multi-channel biomimetic ICMS yielded substantially better compliance discrimination performance than did single-channel linear feedback (25% vs 7.5% error rates for linear and biomimetic, respectively, Figure 6B). We conclude that multi-channel biomimetic ICMS conveys more finely graded force information in both open- and closed-loop contexts.

**Figure 6.**
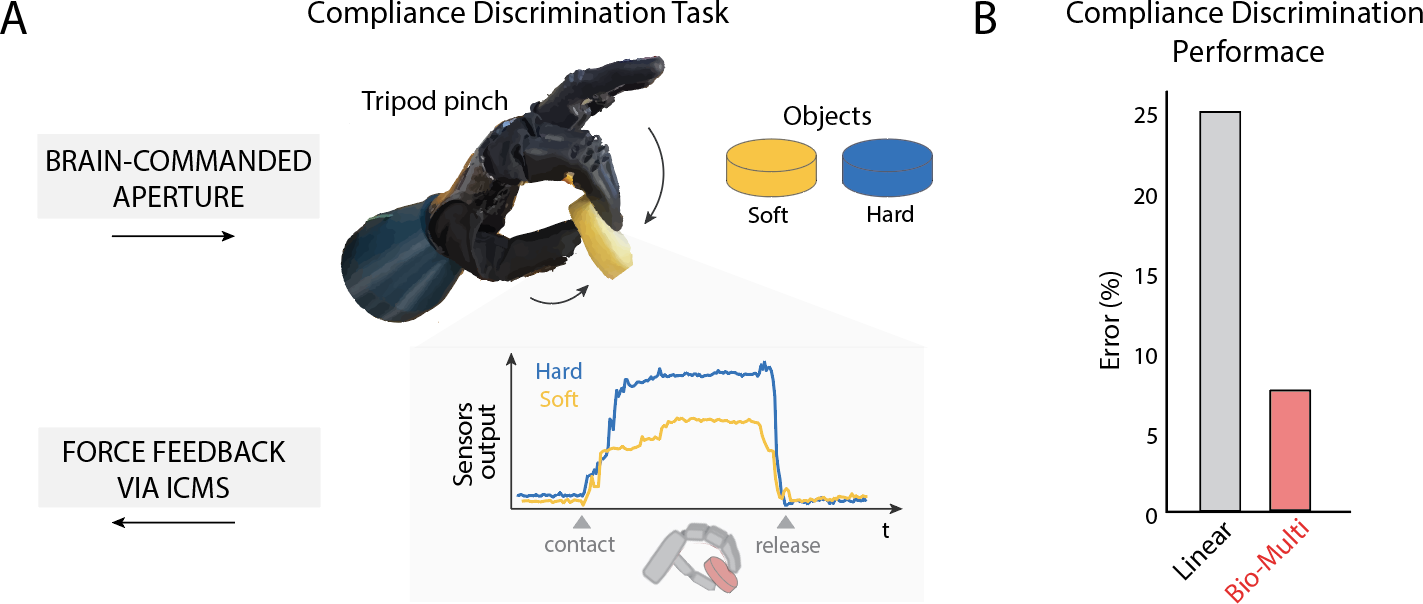
Multi-channel biomimetic feedback is more precise than linear feedback. **A|** Schematic of the Compliance discrimination task using a sensorized bionic hand. **B|** Discrimination performance with linear single-channel and biomimetic multi-channel feedback.

## Discussion

### Discrimination of ICMS amplitude tends to follow Weber’s law

Weber’s law – which states that the JND is proportional to the standard *(17)* – is ubiquitous across sensory modalities. In previous studies with monkeys *(10)* and humans *(6)*, we found that the discrimination of amplitude did not follow Weber’s law, a phenomenon we hypothesized might reflect differences in the trial-to-trial variability of natural sensory responses and otheir ICMS-evoked counterparts *(10)*. In the present study, we find that, on average, JNDs increased in proportion to the standard, consistent with Weber’s law and with natural touch. At the level of single electrodes, however, Weber’s law is often violated, with some JNDs actually lower for the higher standard. The narrow range of perceptible and safe ICMS amplitudes precludes a more systematic investigation of Weber’s law with ICMS. Note that the computed number of discriminable levels is similar with and without the assumption of Weber’s law: assuming Weber’s law, the JNDs are smaller at low amplitudes but larger at high ones than assuming equal JNDs across the range. The large variability across electrodes in their disposition to follow Weber’s law might account for the discrepancy between this and previous studies (along with minor differences in the experimental design).

### Biomimetic ICMS confers higher resolution force feedback

The rationale behind biomimetic feedback is that it evokes more natural patterns of neural activation, which then gives rise to more easily interpretable sensations *(28, 29)*. In experiments with human amputees, biomimetic sensory feedback via peripheral nerve stimulation (PNS) has been shown to improve the function of myoelectric bionic hands *(24, 25)*: With biomimetic feedback, users could more rapidly transfer fragile objects or identify object compliance. With PNS stimulation, however, JNDs were higher with biomimetic feedback, suggesting that the intuitiveness of the biomimetic feedback made up for its lower resolution. Note, however, that this was demonstrated on a single channel and therefore should be replicated before definitive conclusions can be drawn given the channel-dependence of stimulation effects *(24)*. For ICMS-based feedback, biomimetic trains differing in amplitude are even more discriminable than are their amplitude-matched linear trains. The difference between biomimetic and linear ICMS trains is even more pronounced when the stimulation intensity is expressed in terms of overall charge (Supplementary Figure 5B). Whether biomimetic ICMS leads to more intuitive sensations has yet to be systematically investigated, though preliminary findings are promising *(30)*. Regardless, biomimetic ICMS-based feedback offers the additional advantage of higher resolution force information, unlike its PNS counterpart.

### Multi-channel ICMS confers a wider dynamic range of force feedback

Multi-channel ICMS leads to more intense sensations than single-channel ICMS. This finding is perhaps unsurprising given that multi-channel stimulation entails four times more charge delivery than does single-channel stimulation. In fact, multi-channel stimulation less efficiently modulates the overall perceived magnitude compared to single-channel stimulation when expressed as a function of overall charge (Supplementary Figure 6D). The major bottleneck in ICMS-based sensory feedback, however, is that the amplitude used in human experiments is capped at 100 μA, as this level of stimulation has been shown in experiments with monkeys to cause no damage beyond that incurred during implantation *(31)*. Even if this maximum level turns out to be more conservative than it needs to be, evidence suggests that charge per phase is the main determinant of ICMS-induced neuronal damage *(27)*. Accordingly, multi-channel ICMS enables a widening of the dynamic range without increasing the charge per phase on any given electrode.

Beyond widening the dynamic range, multi-channel ICMS improved the resolution of the feedback, as gauged by lower JNDs (Figure 5C). While a reduction in detection thresholds and quickened reaction times with multi-channel stimulation has been previously reported in both humans *(32)* and monkeys *(33, 34)*, the improvement in discriminability has not. In fact, this improvement is inconsistent with the results of experiments with monkeys, in which multi-channel ICMS yielded similar JNDs as did its single-channel counterpart *(35)*. The basis for this discrepancy is unclear.

While JNDs are lower for multi-channel than single-channel ICMS, increasing ICMS amplitude by one JND leads to a greater increment in perceived magnitude with multi-channel than single channel ICMS. If the goal is to evoke a sensation whose intensity is commensurate with the force level, multi-channel ICMS does not confer greater sensitivity to force per se. Rather, the advantage of multi-channel ICMS is that it gives rise to a wider dynamic range of sensations, which can thus signal force over a wider range.

## Conclusion

Biomimetic multi-channel stimulation doubles the dynamic range of the evoked touch sensations, conferring nearly fivefold more discriminable levels of force than does single-channel linear feedback without increasing the charge density.

## Methods

### Participants

This study was conducted under an Investigational Device Exemption from the U.S. Food and Drug Administration and approved by the Institutional Review Boards at the University of Pittsburgh and the University of Chicago. The clinical trial is registered at ClinicalTrials.gov (NCT01894802). Informed consent was obtained before any study procedures were conducted. Participant C1 (m), 57 years old at the time of implant, presented with a C4-level ASIA D spinal cord injury (SCI) that occurred 35 years prior to implant. Participant P2 (m), 28 years old at the time of implant, presented with a C5 motor/C6 sensory ASIA B SCI that occurred 10 years prior to implant. Participant P3 (m), 28 years old at the time of implant, presented with a C6 ASIA B SCI that occurred 12 years prior to implant.

### Cortical implants

We implanted four microelectrode arrays (Blackrock Neurotech, Salt Lake City, UT, USA) in each participant. The two arrays (one medial and one lateral array) in Brodmann’s area 1 of somatosensory cortex were 2.4 mm x 4 mm, with sixty 1.5-mm long electrode shanks wired in a checkerboard pattern such that ICMS could be delivered through 32 electrodes. The two arrays in primary motor cortex were 4 mm x 4 mm, with one-hundred 1.5-mm long electrode shanks wired such that 96 (C1 and P3) or 88 (P2) electrodes could measure neural activity. The inactive shanks were located at the corners of these arrays. Two percutaneous connectors, each connected to one sensory array and one motor array, were fixed to the participant’s skull. We targeted array placement based on functional neuroimaging (fMRI) of the participants attempting to make movements of the hand and arm (all participants) and imagining feeling sensations on their fingertips (participant P2), within the constraints of anatomical features such as blood vessels and cortical topography.

### Intracortical microstimulation (ICMS)

Stimulation was delivered via a CereStim 96 (Blackrock Neurotech). Each stimulating pulse consisted of a 200-µs cathodic phase followed by a half-amplitude 400-µs anodic phase (to maintain charge balance), the two phases separated by 100 µs.

### Multi-channel ICMS

We selected groups of 4 electrodes, referred to as quads. In most cases, the four electrodes had overlapping projected fields (the patch of skin over which the ICMS-evoked sensation is experienced). In one case, one pair of electrodes in the quad had PFs on one digit and the other pair had PFs on the adjacent digit. When stimulating through a quad, all electrodes delivered the same ICMS pulse train synchronously. During each experimental block, we randomly interleaved stimulation through each channel in each quad with stimulation through the entire quad.

### Mechanical skin indentation

Mechanical indentations were delivered with a V-308 voice coil (Physik Instrumente, USA, MA) that drove a probe (tip diameter = 5 mm). Indentations lasted 1 sec, including 0.1-sec on- and off-ramps, matching the temporal profile of the ICMS, and ranged in amplitude from 0.02 mm to 2 mm. The tip was centered on the location of the projected field of an electrode and pre-indented into the skin to ensure maintained contact throughout the experimental block. Mechanical indentations were delivered to participant C1 and to five able-bodied participants (all male, 24-33 years of age) under a separate IRB protocol approved by the University of Chicago.

### Projected fields

Projected fields were collected over multiple years for each electrode and participant. On each trial, a 60-µA, 100-Hz ICMS train was delivered through a given electrode and the participant drew the spatial extent of the sensation on a digital representation of his hand (such as that shown in Figure 1B). The region enclosed by the drawn boundary constituted an estimate of the projected field for that electrode on that session. PFs obtained for each electrode were combined across sessions to obtain a time-averaged estimate.

### Detection thresholds

Detection thresholds for ICMS were measured on each electrode in both sensory arrays on a quarterly basis as described previously *(6)*. We either used a 3 up 1 down transformed staircase or the method of constant stimuli, both in a 2-alternative forced choice (AFC) paradigm targeting 50% detection performance.

### Magnitude estimation

The objective of this paradigm was to assess the relationship between stimulus amplitude and sensory magnitude. On each trial, the participant was presented with an ICMS or a mechanical stimulus and rated its sensory magnitude on a scale of his choosing. If the stimulus was imperceptible, the participant ascribed to it a rating of zero. If one stimulus was twice as intense as another, it was ascribed a rating that was twice as large. The participant was encouraged to use fractions or decimals. At the beginning of each set, the participant was presented with each of the test stimuli in a random order to familiarize them with the stimulus range, as is typically done. The amplitude of the ICMS stimulus varied between 20 and 80 µA while the mechanical stimuli ranged from 0.05 to 2 mm. In all cases the mechanical stimulus was delivered to the location of the projected field for the given electrode. Both ICMS and mechanical stimuli were interleaved throughout each block, with a 5-sec inter-trial interval. For each electrode, each rating was divided by the grand mean of all ratings in the experimental block, then averaged across repeated presentations of each stimulus.

In experimental blocks involving both skin indentations and ICMS, the range of indentation depths was selected in preliminary measurements to match the range of intensities of the electrical stimuli (based on participant ratings) to maximize the overlap and minimize range-related biases *(21)*. To this end, we estimated, at the start of the session and with the participants help, the range of mechanical indentations that evoked sensations of comparable intensity as their ICMS counterparts (which varied from channel to channel). When multiple channels were tested, we selected the weakest and strongest sensations across channels. We then evenly sampled intermediate indentation depths between these two extremes. In some cases, the intensity of the multi-channel ICMS exceeded the maximum indentation that could be delivered with the indenter, precluding comparisons at higher amplitudes.

A minimum of 8 blocks were completed for each channel and condition. Blocks were sometimes distributed across several days to minimize the effects of adaptation *(36)* and maintain participant engagement.

### Amplitude and force discrimination

Participants performed an amplitude discrimination task in a two-alternative forced choice paradigm. On each trial, a pair of stimuli, each lasting 1 sec, was presented with a 1-sec inter-stimulus interval and the participant reported which stimulus was stronger. The standard stimulus, consistent across the experimental block, was paired with a comparison stimulus whose amplitude varied from trial to trial and spanned the standard amplitude. The order of presentation of the standard and comparison stimuli was randomized and counterbalanced. Data were obtained from each electrode over a minimum of 8 experimental blocks, each consisting of 2 presentations (one for each order) of each stimulus pair. The frequency of the ICMS stimuli was either 50 Hz (Figure 3) or 100 Hz (all other experiments).

In force discrimination experiments, trapezoidal force traces (1-sec in duration, 0.1-sec ramps) were converted to ICMS according to a linear or biomimetic mapping. For the linear mapping, the ICMS profile matched that of the force profile, with a peak amplitude proportional to the peak force. For the biomimetic mapping, ICMS consisted of 0.1-sec transients during the on- and off-force ramps whose amplitude matched the peak amplitude of the corresponding linear train and a sustained phase, during which the amplitude was either 30 μA less than the peak amplitude or the detection threshold, whichever was highest. The envelope of the resulting biomimetic trains mimics the envelope of the spiking response to an indentation in S1 *(11)* . On any given experimental block, the standard force was mapped to an ICMS amplitude of 60 µA and comparison forces mapped onto amplitudes that ranged from 40 to 80 µA (Figure 4A). Note that the JNDs were the same at 50 and 100 Hz, as has been previously found in experiments with monkeys *(10)*.

To test whether ICMS amplitude discrimination was subject to Weber’s law, we selected two standards – either at 50 and 70 μA – and paired them with comparison amplitudes that spanned a range of ±16 µA around the standard (based on previously observed JNDs). Tests for both standards were interleaved and randomized within an experimental block. This design contrasts with that adopted in previous experiments testing Weber’s law, in which the two standards were at the extremes and comparisons fell between the extremes *(6, 10)*. The narrow range of the standards was adopted such that none of the comparisons fell below threshold or above the safety limit of 100 μA.

The same behavioral paradigm was used to measure amplitude JNDs with the mechanical indenter as for ICMS. The indentation depths varied between 0.85 and 1.15 mm with a standard amplitude of 1 mm and an onset- and offset-ramp speed of 5 mm/s.

#### Analysis

### Equal intensity contours

We fit a power function to the magnitude ratings, *M_m_*, of the skin indentations:

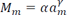

where *a_m_* is the amplitude, and α and γ are fitted parameters. We fit a linear function to the magnitude ratings, *M_e_*, of the ICMS:

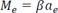

where *a_e_* is the ICMS amplitude We then derived an equal intensity contour by finding the conditions under which *M_m_* equals *M_e_:*

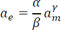

### Force approximation

To convert skin indentations into equivalent forces, we measured the relationship between skin indentation and exerted force using a Universal Testing Machine (Instron, Supplementary Figure 8). Data was collected from 15 individuals (8 women, 7 men) aged between 21 and 30. In brief, we indented the skin over a range of depths and measured the force required to achieve that indentation. We then used the resulting relationship to estimate the forces applied to the skin by the indenter in the magnitude estimation task.

### Psychometric functions

Psychometric functions were fit with a logistic function:

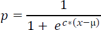

Where *p* is the probability of judging the comparison stimulus as more intense than the standard, *x* is the amplitude, *c* is the slope, and *µ* the point of subjective equality (PSE). The just noticeable difference (JND) is half the difference between the amplitudes that yield a *p* of 0.25 and 0.75.

### Discriminable levels

To estimate the number of discriminable levels for ICMS we used the formula:

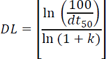

where DL is the number of discriminable levels, *dt_50_* the 50% detection threshold, and k the weber fraction for that channel, respectively. For touch we estimated the number of levels based on a computed Weber fraction of 1.083 and *dt_50_* of 0.05 mm.

### Decoding of hand aperture

Neural signals from M1 arrays were recorded at 30 kHz using the NeuroPort system (Blackrock Neurotech, Salt Lake City, UT, USA). The data were high-pass filtered with a 1^st^ order Chebychev filter above 750 Hz. Whenever the signal crossed a threshold (−4.5 RMS, set at the start of each recording session), a spiking event was recorded. Spikes were binned in 20-ms bins for decoding, then smoothed with an exponential filter of 440ms. Only channels with firing rates above 0.5Hz were used.

To decode aperture, we used a recurrent neural network that consisted of one middle layer of 150 long short-term memory (LSTM) nodes implemented online in subject C1. Network hyper-parameters network were optimized to achieve the best cross validated fit. For each session, we first harnessed M1 responses as the participant attempted to grasp an object in virtual reality, following the motion of a virtual hand. During training, contact with the object triggered ICMS (linear single-channel and biomimetic multi-channel) to expose the decoder to the activity in M1 evoked by ICMS delivered to S1, which would otherwise have disrupted decoder performance when not accounted for *(37)*. Offline training lasted less than 5 minutes. An LSTM decoder of hand aperture was constructed based on these data, the participant proceeded to use the decoder to perform the compliance discrimination task described below. Real time inference was performed with a lag shorter than 10 ms.

### Compliance discrimination task

On each trial, the participant was instructed to squeeze two foam pucks of differing compliance with the bionic index and thumb. The output of the sensor on the thumb and index finger were summed separately to yield two force signals, one stemming from each digit, which then drove ICMS delivered through electrodes with PFs on the corresponding digit. These sensor signals were converted into ICMS using either a linear or biomimetic algorithm. For the linear algorithm, ICMS amplitude was proportional to (summed) sensor output, F(t):

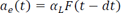

where *dt* is the bin size (20 ms) and *α_L_* is a parameter, chosen such that the ICMS amplitude spanned the range from 20-90 µA based on preliminary measurements of the sensor output ranges in this task.

For the biomimetic algorithm, ICMS amplitude was proportional to both the sensor output and its derivative:

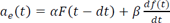

For this, α and β were selected such that the resulting ICMS trains peaked at comparable amplitudes as with the linear feedback, based on the aforementioned preliminary testing. For the biomimetic feedback,

ICMS was delivered through quads of electrodes with PFs on the thumb and index, with the same stimulus applied to all 4 channels.

## Acknowledgments

We would like to thank the participants for their generous contribution to the advancement of science. This work was supported by NINDS grants UH3 NS107714 and R35 NS122333.

## Disclosures

NH and RG serve as consultants for Blackrock Neurotech, Inc. RG is also on the scientific advisory board of Neurowired LLC. MB, JC, and RG receive research funding from Blackrock Neurotech, Inc. though that funding did not support the work presented here.

## Supplementary Figures

**Supplementary Figure 1.**
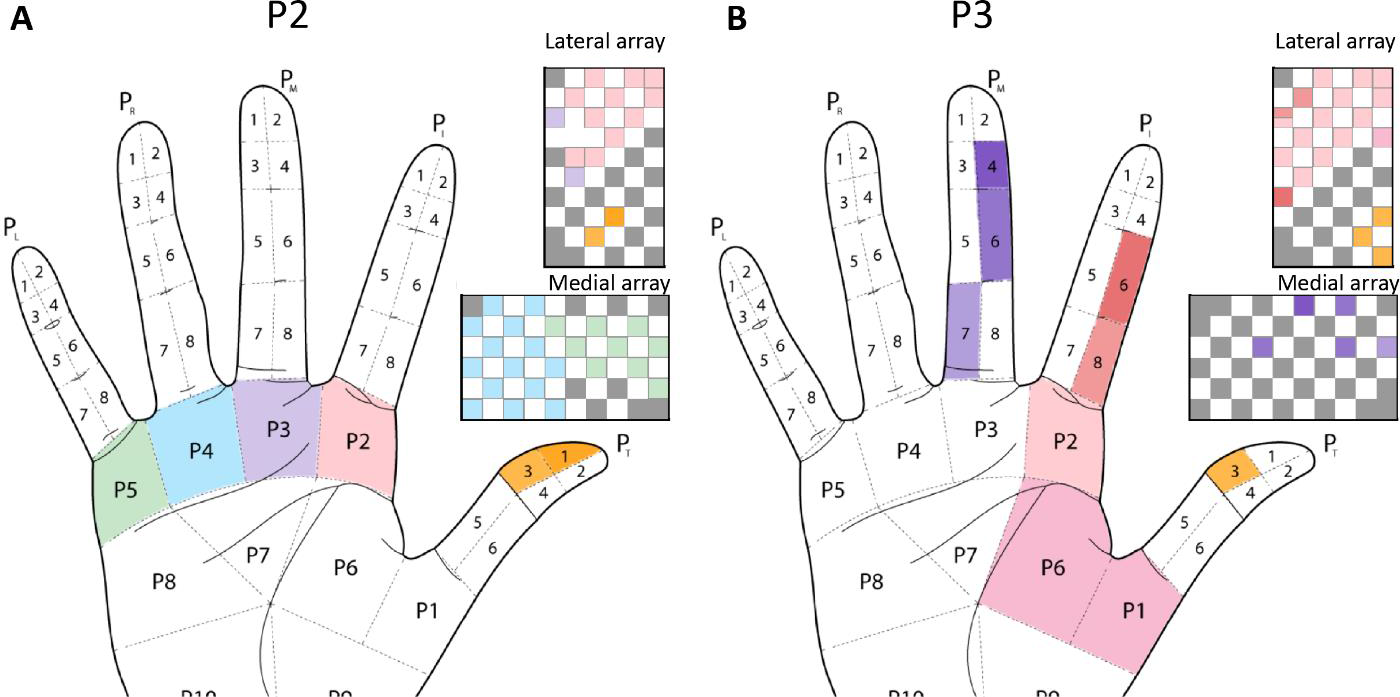
**A|** Locations of projected fields for each S1 channel for participant P2. **B|** Locations of projected fields for each S1 channel for participant P3. The top array is medial, bottom one lateral. Colors denote the location of the projected field. Gray squares denote electrodes that evoked sensations on the dorsum of the hand, and white squares denote unwired electrodes.

**Supplementary Figure 2.**
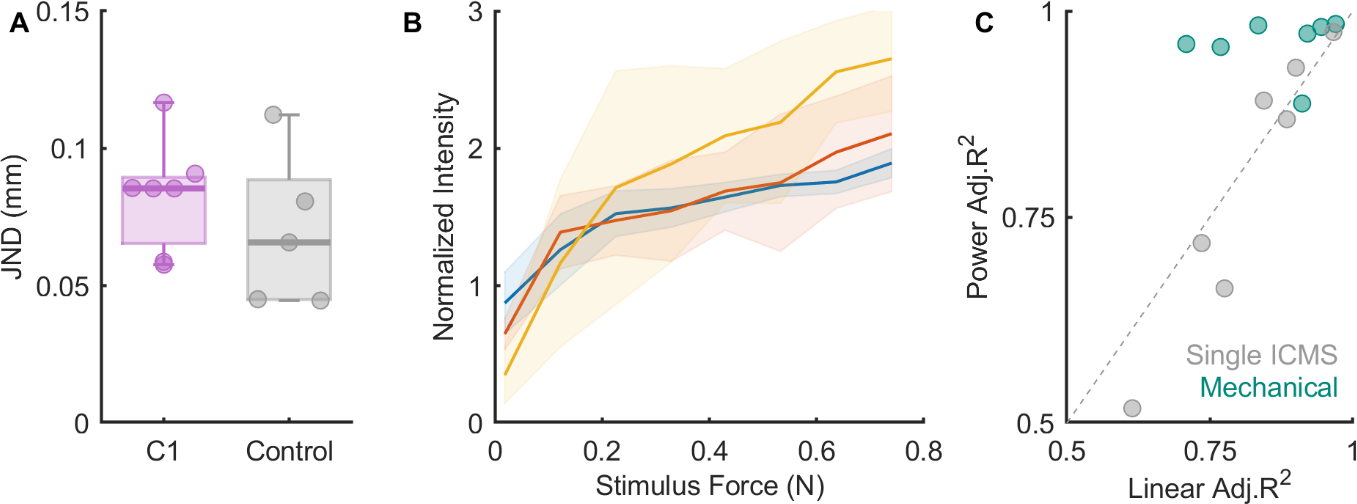
**A|** Mechanical JNDs for participant C1 (each point denotes performance on an individual digit) and for 5 able-bodied controls (each point denotes performance of one participant on the tip of the middle finger). The participant’s ability to distinguish the depth of indentation is comparable to that of the controls. **B|** Normalized intensity ratings from participant C1 on 3 different digits (denoted by different colors). Ratings are consistent across digits. Note that the normalization was performed based on the grand mean rating, which included ratings of single-channel ICMS stimuli and tended to be weaker than the mechanical ones. **C|** Goodness of fit for ICMS or mechanical intensity ratings when using power vs. linear functions. Power functions provide better fits for intensity ratings of skin indentations but not ICMS.

**Supplementary Figure 3.**
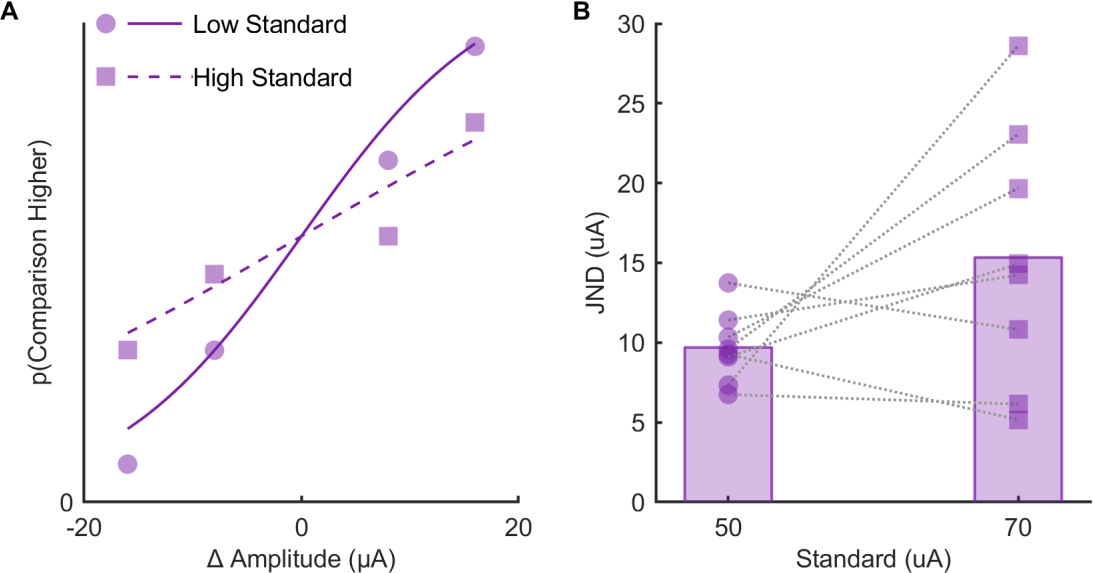
**A|** Example psychometric functions with two standards for one electrode. **B|** JNDs with 2 standards for 8 electrodes. While mean JNDs were higher for the higher standard, the effect was not significant (paired t-test, t = 1.8, p = 0.11).

**Supplementary Figure 4.**
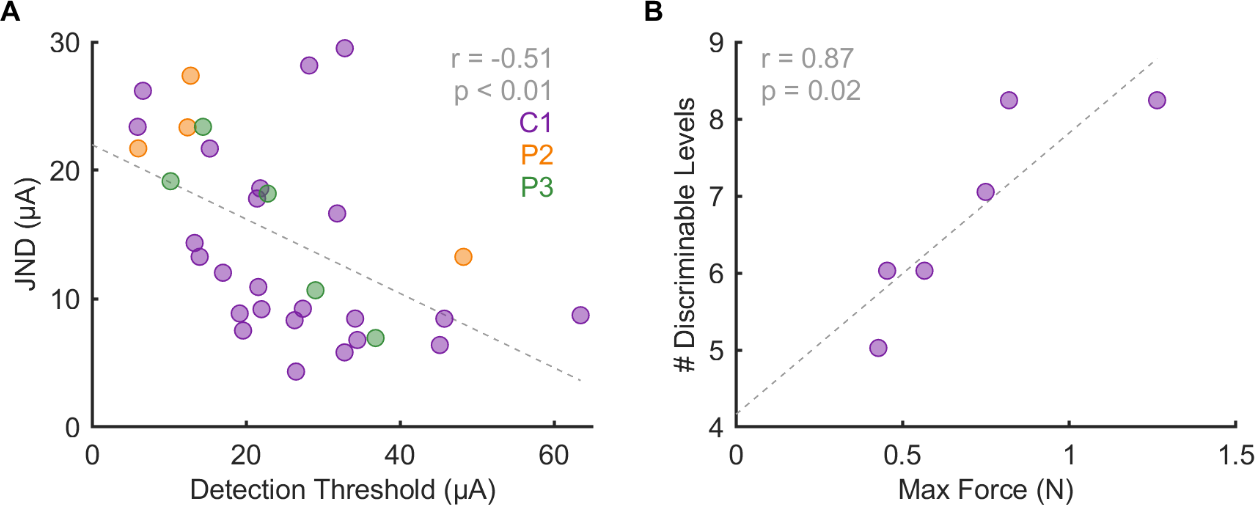
**A|** The discriminability of an electrode (JND) is inversely correlated with its detection threshold. **B|** Electrodes that had a broader dynamic range (i.e., a greater maximum equivalent force) tended to yield more discriminable levels.

**Supplementary Figure 5.**
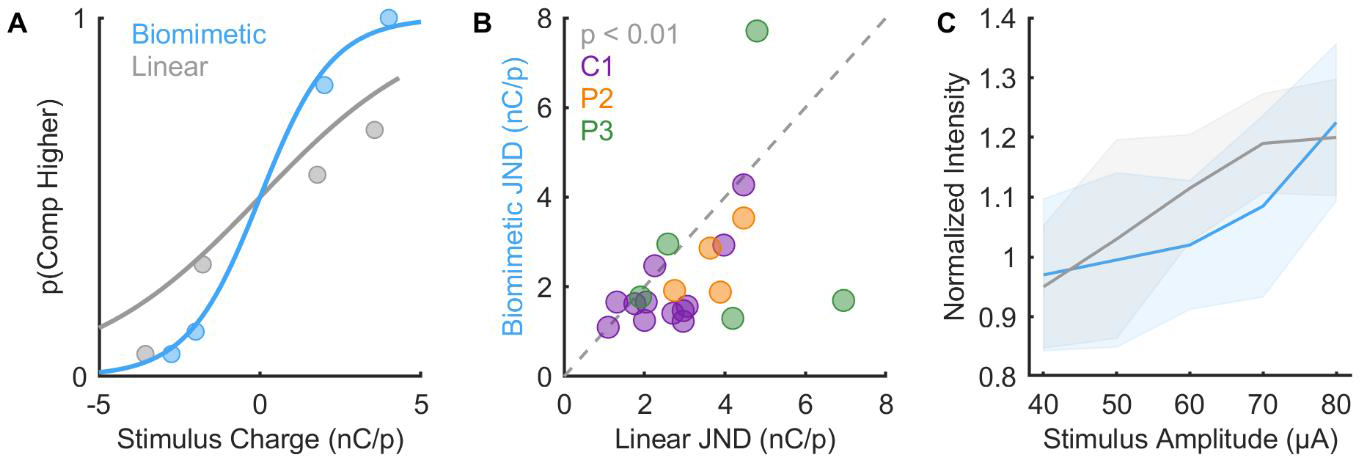
Discrimination performance expressed in terms of injected charge. **A|** Psychometric functions for biomimetic (blue) and linear (gray) trains, expressed in terms of mean charge per phase, for one electrode. **B|** JNDs for biomimetic and linear stimuli across participants and electrodes (n = 20). Biomimetic stimuli yield significantly lower JNDs than linear stimuli, expressed in terms of charge per phase (Wilcoxon signed-rank test, Z = 2.8, p < 0.01). **C|** Normalized intensity ratings for one channel. Linear ICMS (gray) does not give rise to significantly more intense sensations than does biomimetic ICMS (blue) when comparing stimuli with the same maximum amplitude. The mean relative intensity of the biomimetic stimuli was 92 ± 3% of the intensity of the linear ones (n = 5 electrode).

**Supplementary Figure 6.**
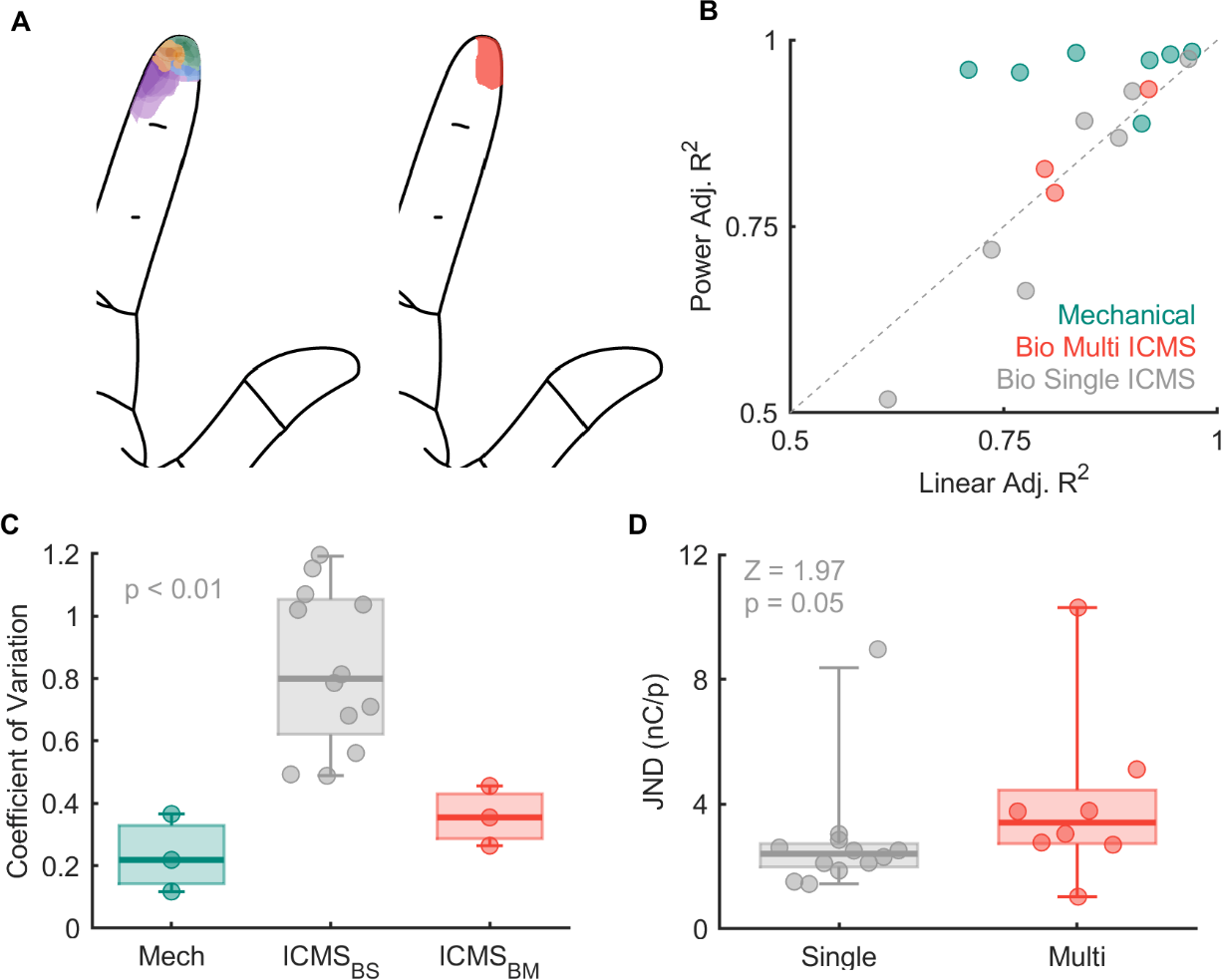
**A|** Projected field locations for electrodes stimulated individually (left) and simultaneously (right) in one quad from participant C1. **B|** Linear fits are equivalent to power law fits for single-channel of multi-channel ICMS but power law fits are better for mechanical stimuli. **C|** Variability of magnitude ratings is lower for multi-channel stimulation than for single-channel stimulation (Tukey’s post-hoc test, p < 0.01) and statistically equivalent to its mechanical counterpart (p = 0.45). **D|** JNDs are significantly lower for single than multi-channel biomimetic stimulation when expressed in terms of charge per phase (1-way ANOVA, F = 36.1, p < 0.01).

**Supplementary Figure 7.**
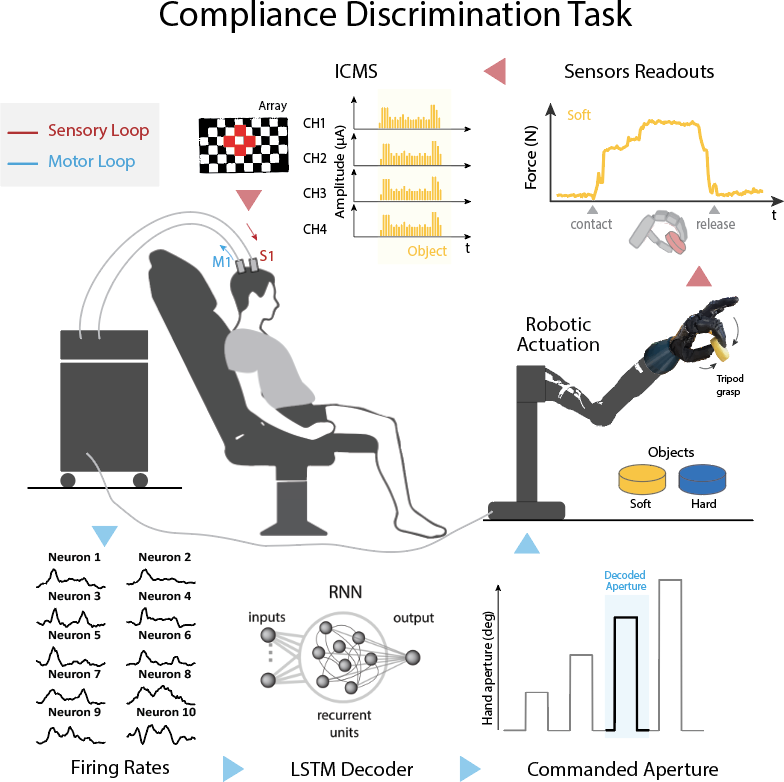
Pipeline for the compliance discrimination task. At trial start, the subject attempts to grasp a foam puck. An LSTM decoder infers the attempted aperture based on signals from M1. Upon object contact, the output of sensors on the thumb and index finger drives ICMS according to a single-channel linear algorithm or a multi-channel biomimetic one. Linear feedback is delivered through a single electrode, biomimetic feedback through a quad with overlapping PFs.

**Supplementary Figure 8.**
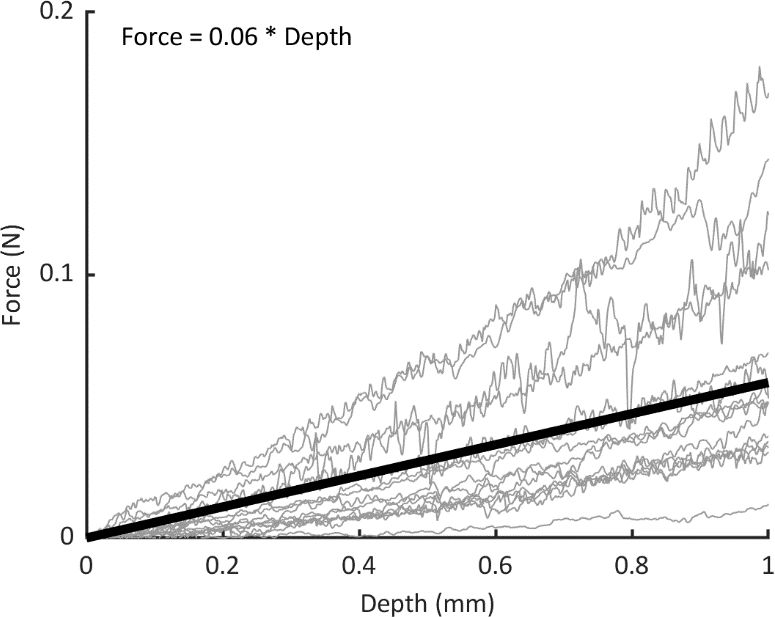
Indentation depth vs force for 15 human participants using a 2-mm diameter tip with a universal testing machine (Instron). Each line denotes data from one participant, thick black line denotes the mean.

## Notes

### Competing Interest Statement

NH and RG serve as consultants for Blackrock Neurotech, Inc. RG is also on the scientific advisory boards of Braingrade GmbH and Neurowired LLC. MB, JC, and RG receive research funding from Blackrock Neurotech, Inc. though that funding did not support the work presented here.

### Summary of Updates

Revisions throughout. Addition of a closed loop task with a bionic hand to demonstrate real world use.

